# Role of Water Models in Simulations of Ion Conduction in Potassium Channels

**DOI:** 10.1101/2025.10.24.684299

**Authors:** Stefano Bosio, Diego Gazzoni, Carmen Domene, Matteo Masetti, Simone Furini

**Author notes:** Corresponding authors: Matteo Masetti,; Simone Furini.

## Abstract

Potassium channels exhibit high selectivity and conductance, yet the atomic details of ion permeation, particularly the involvement of water molecules, remain debated. Two main conduction mechanisms have been proposed: the hard knock-on, in which ions traverse the selectivity filter in direct contact, and the soft knock-on, which involves co-permeation of water molecules. Using microsecond molecular dynamics simulations with the OPC water model, the AMBER19SB protein force field, and the 12-6-4 Sengupta et al. ion model, we observed that both hard and soft knock-on mechanisms are accessible and, notably, can reversibly transition in the MthK and KcsA channels across all simulated membrane potentials. These reversible transitions contrast with previous observations using the TIP3P water model, where water entry either disrupted conduction or was expelled, favoring exclusive hard knock-on events. Our results suggest that the choice of the water model, force field, and ion parameters significantly influences the observed conduction mechanism. Importantly, the coexistence of hard and soft knock-on in these simulations provides a potential reconciliation between structural data supporting hard knock-on and streaming potential measurements demonstrating water co-permeation. These findings reintroduce soft knock-on as a viable conduction mechanism and highlight the critical role of simulation parameters in reproducing potassium channel permeation behavior.

## Introduction

Potassium channels (K^+^ channels) are transmembrane proteins that facilitate the selective transport of potassium ions across biological membranes down their electrochemical gradient.(*1*) These channels are widely expressed in both excitable and non-excitable cells across virtually all living organisms,(*2*) and play critical roles in fundamental cellular processes, such as electrical signalling, osmoregulation, and maintaining ionic balance.(*3*) Owing to their central physiological role, dysregulation of K^+^ channels has been implicated in a broad range of diseases, including cardiac arrhythmias,(*4*) neurological disorders,(*5*) and various forms of cancer.(*6*) Despite considerable sequence and functional diversity, K^+^ channels share a conserved architecture. The pore-forming domain typically assembles as a tetramer (or a pseudo-tetramer in the case of the K2P family) with subunits arranged in a fourfold-like symmetry around the pore axis. The ion permeation pathway is formed by two transmembrane (TM) helices contributed by each subunit, which are connected by a pore helix and a highly conserved selectivity filter (SF), containing the TVGYG sequence (**Figure 1A**).(*7*) The SF contains five consecutive K^+^ coordination sites, designated S0 to S4 from the extracellular side. At these sites, potassium ions are coordinated by backbone carbonyl oxygen atoms of the protein, with the exception of S4, where side-chain oxygens from a highly conserved threonine residue contribute to the coordination shell. On the intracellular side of S4, the pore widens into an intracellular cavity, C, which, in the open state, is continuous with the intracellular compartment (**Figure 1B**).

**Figure 1.**
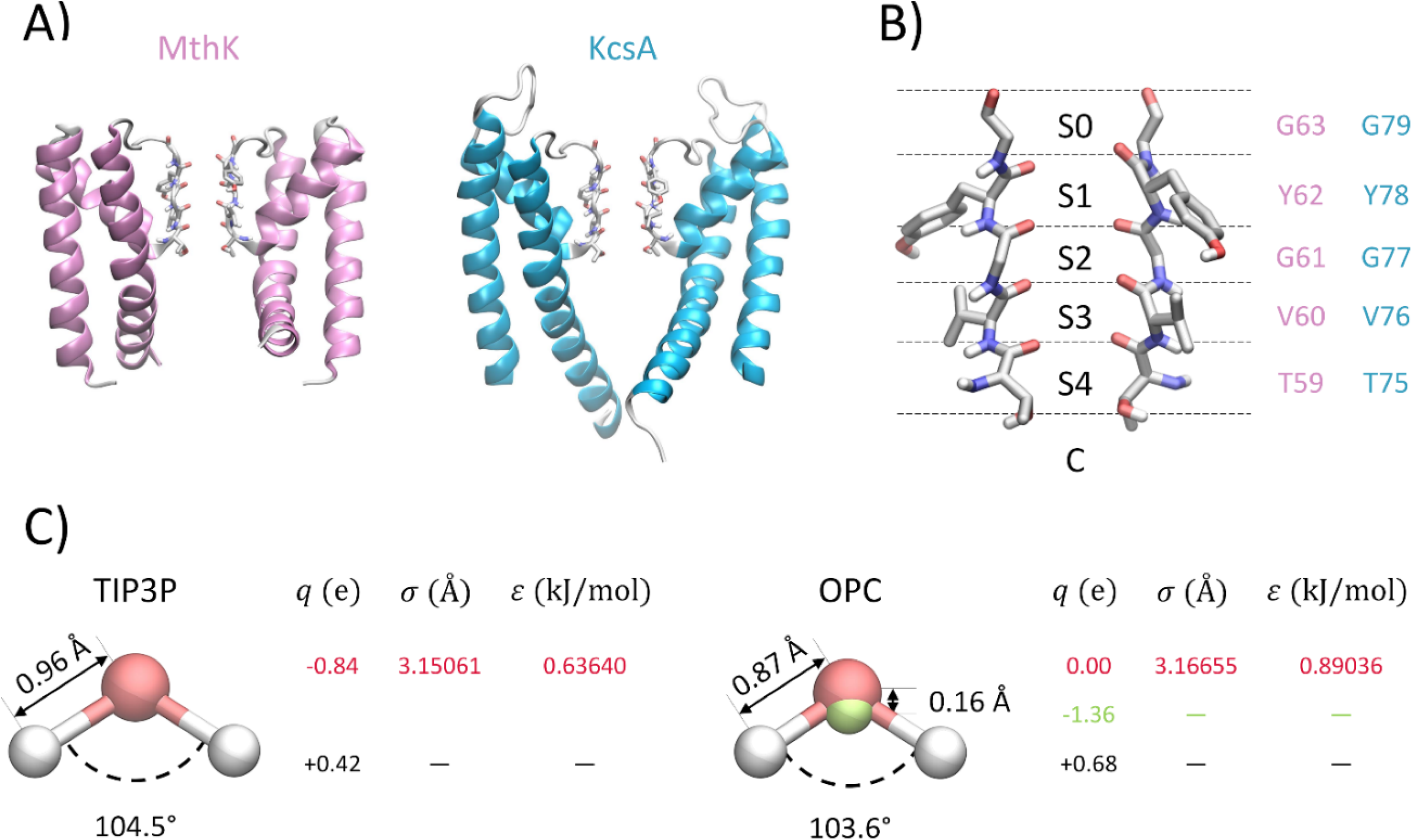
Ion channels and water models. (A) Structures of the pore domains of KcsA (left) and MthK channels (right). Only two opposing subunits are shown for clarity. The selectivity filter (SF) is depicted in licorice representation. (B) Close-up view of the SF, highlighting the five potassium ion binding sites, S0-S4, and the cavity, C. Residue numbering is shown in pink for MthK and light blue for KcsA. (C) Internal coordinates and non-bonded parameters of the TIP3P and OPC water models.

Molecular Dynamics (MD) simulations and related methods have significantly improved our understanding of K^+^ channel function by complementing experimental structural data with time-resolved information.(*8*) In particular, these approaches have been instrumental in identifying key energetic and structural determinants underlying ion selectivity and conduction,(*9, 10*) as well as conformational changes involved in channel gating and ion permeation.(*11, 12*) Despite these advances, a comprehensive and quantitatively accurate picture of ion permeation through K^+^ channels remains elusive.(*13*) For example, conductance values estimated from simulations for K+ channels are typically an order of magnitude lower than those measured experimentally.(*14–16*) Furthermore, the mechanistic features of the conduction process inferred from simulations, including the occupancy and dynamics of ions within the SF, are highly sensitive to the simulation setup and the choice of force field, leading to controversial interpretations of experimental observations.(*17*) Specifically, two main permeation models have emerged: the “soft knock-on” model, in which K^+^ ions permeate with interposed water molecules in a 1:1 ratio,(*18, 19*) and the “hard knock-on” model, in which ions move by direct ion–ion contact without intervening water molecules.(*20, 21*) Experimental data from 2D-IR spectroscopy,(*22*) streaming potentials,(*23*) and early free-energy calculations(*24, 25*) supported the soft knock-on mechanism. However, 2D-IR measurements were later shown to be compatible with hard knock-on conduction,(*14*) which is also supported by experimental X-ray crystallographic structures obtained in the presence of electric field stimulated conduction.(*26*)

A key limitation of common biomolecular force fields, widely recognized as a source of discrepancies between simulations and experiments, is the lack of explicit polarization.(*27*) In the context of ion channels, this issue has been addressed by Jing et al., who investigated the relative stability of distinct occupancy states of KcsA SF using both the AMOBEA polarizable force field and CHARMM36m with Electronic Continuum Correction (ECC).(*28*) The latter provides a general framework for capturing effective polarization in non-polarizable force fields by rescaling the charges of ionized groups and ions by a factor 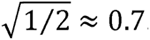. More recently, Hui et al. expanded this investigation by studying ion conduction in three potassium channels, combining the ECC approach with the CHARMM36m and Amber14SB force fields, and systematically testing various scaling factors between 1.0 and 0.65.(*29*) Despite these efforts to capture polarization effects more accurately, other factors in the simulation setup can also influence the behaviour of ion channels and should be carefully considered. In this respect, we previously demonstrated that Amber14SB better preserves the conductive state of the SF over microsecond-long simulations than CHARMM36m.(*15, 30*) However, to the best of our knowledge, the potential impact of water models on K^+^ ion conduction has so far received little attention.

By directly interacting with ion channels and the ions that are transported, water molecules play an active role in channel function, influencing hydration, electrostatic screening, and the energetics of ion conduction and gating. The TIP3P water model (**Figure 1C**),(*31*) widely used for its compatibility with different families of force fields and its computational efficiency, is increasingly recognized as suboptimal for accurately capturing key water properties. For instance, while TIP3P reproduces bulk properties such as the enthalpy of vaporization in reasonable agreement with experiments, it underestimates the density, overestimates the dielectric constant, and fails to reproduce dynamic properties such as the self-diffusion coefficient.(*32, 33*) Among rigid and non-polarizable water models, recent studies highlight the four-point OPC water model (**Figure 1C**)(*34*) as a more accurate alternative, offering enhanced accuracy in hydration properties of proteins and nucleic acids, as well as in describing ion-water interactions, in better agreement with experimental data.(*35, 36*)

In this work, we compare the conduction properties of two prototypical potassium channels, KcsA and MthK, using the TIP3P and OPC water models under different applied voltages. We highlight that simulations with TIP3P employed the Amber14SB force field(*37*) for the protein, and Joung-Cheatham parameters(*38*) for the ions. In contrast, simulations with OPC adopted Amber19SB,(*39*) for which OPC is the recommended choice, and the 12-6-4 Sengupta et al. ion model,(*40*) which is specifically developed for use with the OPC water model. The main finding was that simulations using OPC exhibited both hard and soft knock-on mechanisms of conduction coexisting, whereas simulations using TIP3P showed that hard knock-on was the exclusive, conduction mechanism. The coexistence of soft and hard knock-on conduction in simulations with OPC was observed for both ion channels, KcsA and MthK, and across all the simulated membrane potentials (100, 200, and 400 mV). These findings demonstrate the critical influence of water model choice in dissecting the atomic mechanisms of conduction in K^+^ channels and in addressing the long-standing debate between soft and hard knock-on conduction.

## Methods

### Molecular Dynamics simulations

Atomic coordinates for the KcsA channel with the Glu71 to Ala mutation were obtained from its experimentally determined open/conductive state (Protein Data Bank entry 5VK6).(*41*) The MthK channel coordinates were taken from PDB entry 3LDC.(*42*) In both cases, the entire transmembrane domain of the channels was included in the models, spanning residues Trp26 to Gln121 for KcsA, and Val18 to Ile99 for MthK. Model systems were assembled using the CHARMM-GUI web server.(*43*) Channel orientations within the lipid bilayer followed those specified by the Orientations of Proteins in Membranes (OPM) database.(*44*) KcsA was embedded in a 3:1 mixture of 1-palmitoyl-2-oleoyl-glycero-3-phosphocholine (POPC) and 1-palmitoyl-2-oleoyl-sn-glycero-3-phosphate (POPA), while MthK was simulated in a pure POPC membrane, consistent with previous MD studies of this channel. Differences in lipid composition were not expected to significantly influence ion conduction behaviour on the simulated timescales.(*45*)

Two simulation setups were used for each channel, differing in water models and force field parameters for protein and ions. Systems employing the TIP3P water model(*31*) used the Amber14SB force field(*37*) along with Joung-Cheatham ion parameters.(*38*) Conversely, systems using the OPC water model(*35*) applied the Amber19SB force field(*39*) combined with the 12-6-4 ion model of Sengupta et al.(*40*) Each system contained 150 mM KCl, with potassium ions manually placed at S4, S2, and S0 sites of the SF. van der Waals interactions were truncated at 9 Å, and standard AMBER 1-4 scaling was used. For each combination of simulation setup and channel, simulations were performed under applied membrane potentials of +100 mV, +200 mV, and +400 mV, with multiple independent replicas run for each condition to ensure statistical robustness.

System equilibration followed a multi-step protocol (1, 2, 2, 5, 10, 30, and 100 ns) with harmonic restraints applied along the membrane normal (z-axis) to the centre of mass of lipid headgroups and to the root mean square deviation (RMSD) of the protein backbone and side-chain heavy atoms relative to the starting configuration. Force constants were initially set at 1000, 4000, and 2000 kJ mol^-1^ nm^-2^ for lipid headgroups, backbone atoms, and side chain atoms, respectively, and were progressively reduced through 400, 2000, 1000; 400, 1000, 500; 200, 500, 200; 40, 200, 50; 0, 50, 0; and, finally 0, 0, 0 kJ mol^-2^ nm^-2^. A timestep of 1 fs was used for the first three equilibration stages and 2 fs for the remaining steps. Long-range electrostatics were calculated using the Particle Mesh Ewald (PME) method with a grid spacing of 1.0 Å.(*46, 47*) Bonds involving hydrogen atoms were constrained using the LINCS algorithm(*48, 49*) for the protein and SETTLE for water molecules.(*50*) Temperature was maintained at 310.15 K through the v-rescale thermostat(*51*) with a coupling constant of 1.0 ps-1. During equilibration, pressure was kept at 1 atm with a C-rescale barostat and a damping constant of 5 ps.(*52*) Membrane potentials were simulated by applying a constant electric field along the axis perpendicular to the lipid bilayer. Simulations under applied electric field were conducted in the NVT ensemble as described previously.(*53, 54*). Simulations were performed using Gromacs2023(*55*) with the exception of simulations of KcsA with the TIP3P water model, which were previously generated using NAMD2.12(*56*) as described in (*16*). The cumulative simulation time exceeded 200 µs (**Table S1**).

### Markov State Models

Trajectory analyses were performed using MDAnalysis(*57*) together with the SciPy ecosystem.(*58*) Visual Molecular Dynamics (VMD) was used to visualize trajectories and generate molecular representations of the systems.(*59*) Analysis of ion and water occupancy states was performed using a protocol previously established in our earlier work.(*16*) Each frame of every trajectory was converted into a symbolic string describing the occupancy pattern of the cavity, C and of binding sites S0-S4 (**Table S2** and **S3**) for subsequent analyses. This encoding scheme yielded a discrete-state representation of filter configurations, suitable for constructing Markov State Models (MSMs) from the individual trajectories.

MSMs were built for KcsA and MthK using simulation data obtained at all applied membrane potentials (+100 mV, +200 mV, +400 mV). The transition matrix was estimated as:

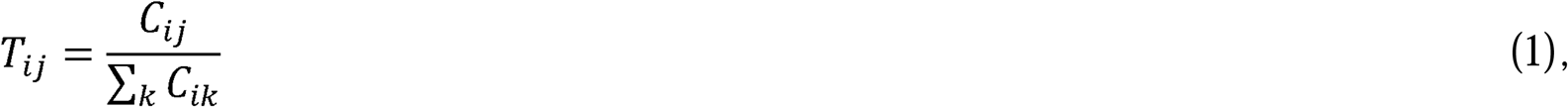

where *T*_*ij*_ is the probability of transitioning from state *j* to state *i*, and *C*_*ij*_ is the number of transitions observed between these states during the entire simulation time. Each MSM transition matrix has a dominant eigenvalue of 1, whose corresponding eigenvector represents the equilibrium distribution.(*60*) All other eigenvalues (*J*_*i*_) have magnitudes less than 1 and are associated with relaxation times:

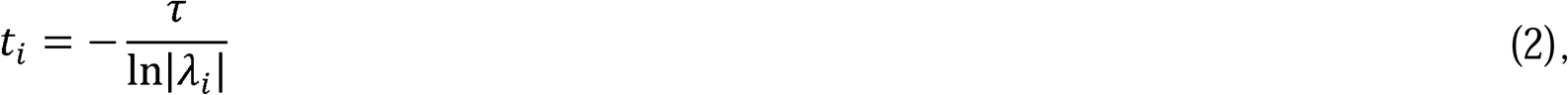

where *T* is the lag time used for sampling. A sampling interval of 1 ns was used for all analyses. Network visualizations of state transitions were generated with Gephi using the ForceAtlas2 algorithm to arrange the nodes.(*60*) Occupancy states were encoded using the regular expression [Kw][Kw-][Kw-][Kw-][Kw-][Kw-]. The first character indicated cavity occupation (“K” = ion, “w” = water molecules without ions). Subsequent characters described the S4-S0 sites (“K” = ion, “w” = water, “– “ = vacant).

## Results

MD simulations of KcsA and MthK channels using the OPC and the TIP3P water models were performed in the presence of an electric field corresponding to membrane potentials of 100, 200, and 400 mV, with multiple replicas for each simulated condition (**Table S3**). When the TIP3P model was used, no water molecules were observed in binding sites S2 and S3 (**Figure 2**), regardless of the channel type or membrane potential. The complete absence of water molecules in S2 and S3 implies that all conduction events follow the hard knock-on mechanism*, i.e.* no water molecules are co-transported with ions. The average conductance, computed by counting the number of conduction events over time, underestimates the experimental value for both channels (**Table 1**).(*61, 62*) The estimated conductance increases significantly with the membrane potential. In detail, the estimated conductance at 400 mV is 6-7 times higher than at 100 mV, highlighting the importance of performing MD simulations at physiological membrane potentials when comparing with experiments. These results of MD simulations using the TIP3P model are consistent with previous observations in the literature.(*14, 16*)

The behaviour of water molecules, and consequently of ions, in the SF changes markedly when the OPC model is adopted. In simulations with OPC, water molecules bind to the SF sites S2 and S3, as revealed by the presence of water density peaks at the core of the SF, which are completely absent in simulations with TIP3P (compare the left and right plots in **Figure 2**). The distribution of water molecules in the intracellular cavity, at the extracellular entrance of the SF, and between the SF and the P-loops is similar in simulations with the two water models, suggesting that other aspects of channel dynamics, such as hydrophobic gating,(*63–65*) are unlikely to be affected by the choice of water model. Conversely, the presence of water molecules inside the SF clearly implies an influence of water on the ion conduction mechanism.

**Table 1.**
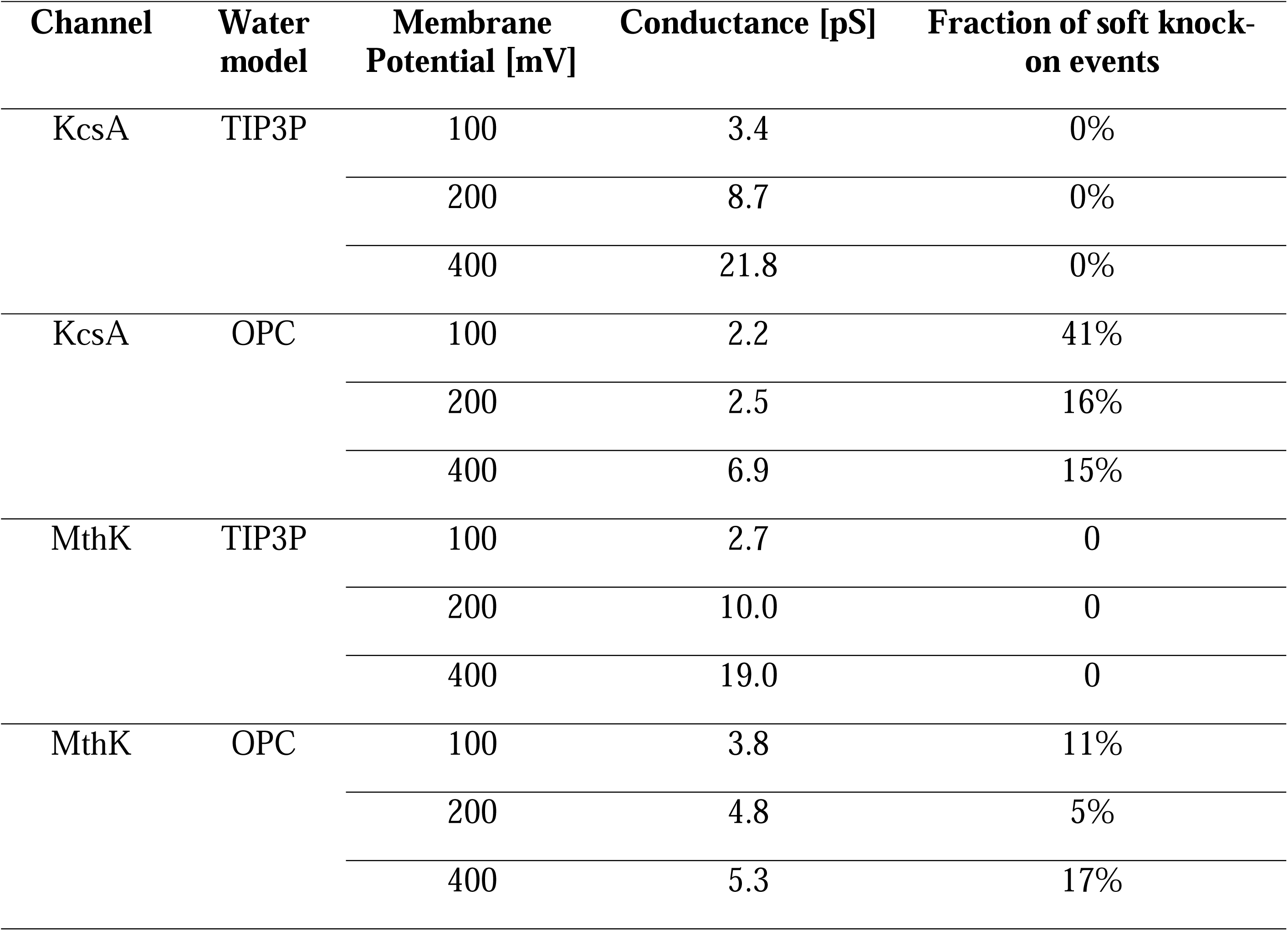
Average properties of conduction in simulations estimated from simulations of KcsA and MthK.

| Channel | Water model | Membrane Potential [mV] | Conductance [pS] | Fraction of soft knock-on events |
| --- | --- | --- | --- | --- |
| KcsA | TIP3P | 100 | 3.4 | 0% |
|  |  | 200 | 8.7 | 0% |
|  |  | 400 | 21.8 | 0% |
| KcsA | OPC | 100 | 2.2 | 41% |
|  |  | 200 | 2.5 | 16% |
|  |  | 400 | 6.9 | 15% |
| MthK | TIP3P | 100 | 2.7 | 0 |
|  |  | 200 | 10.0 | 0 |
|  |  | 400 | 19.0 | 0 |
| MthK | OPC | 100 | 3.8 | 11% |
|  |  | 200 | 4.8 | 5% |
|  |  | 400 | 5.3 | 17% |

**Figure 2.**
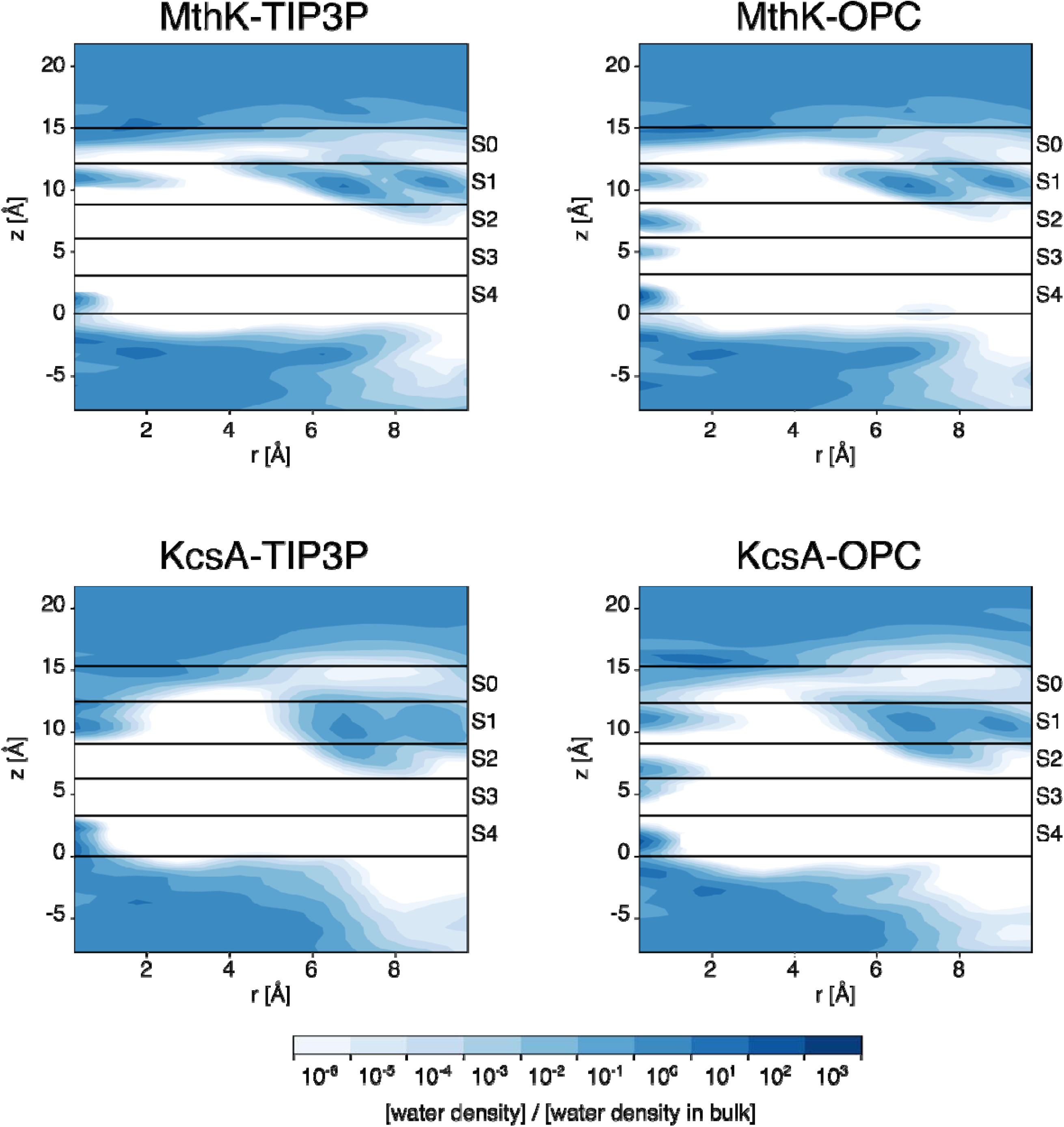
Average density of water molecules. The average density was calculated by combining snapshots from all replicas. The distance from the channel axis, r, was discretised into cylindrical shells spaced 0.5 Å apart, and the axial coordinate was also discretised with a bin size of 0.5 Å. The density was then normalised by the average bulk water density. The boundaries of binding sites S0 to S4, as defined by the average positions of the delimiting oxygen atoms, are shown as black lines.

The entry and exit of water molecules into and from the SF in simulations with the OPC water model is a reversible process, observed multiple times across independent replicas at all the simulated membrane potentials (**Table S3**). As an example, **Figure 3** shows one such event in a simulation of the MthK channel at 200 mV. The entrance of a water molecule in the SF is preceded by the formation of a gap – an empty binding site – in S2, together with the presence of an ion in S3, a water molecule in S4, and an ion in the cavity (state a in **Figure 3**). From this configuration, the chain of ion-water-ion at the intracellular side of the SF moves upward in a sequence of concerted states that finally release an ion and a water molecule in the extracellular compartment (states b-c-d in **Figure 3**). Similar events were observed to occur in simulations at other membrane potentials, and in the KcsA channel, as described in more quantitative terms in the next paragraphs. The estimated conductance at 100 mV is comparable between simulations with TIP3P and OPC and in both cases it is significantly lower than the experimental value (**Table 1**). Consistent with the TIP3P simulations, the conductance increases with increasing membrane potential, however, in the OPC simulations the magnitude of this increase is smaller. The conductance at 400 mV is approximately 1-3 times higher than that at 100 mV in OPC simulations, whereas the corresponding ratio is 6-7 in TIP3P simulations. The major difference between the two water models is the presence of soft knock-on conduction events in OPC simulations which are completely absent in TIP3P simulations. While the hard knock-on mechanism remains the dominant, a significant fraction of soft knock-on events, ranging between 5% and 40%, depending on the channel model and membrane potential, was observed in OPC simulations (**Table 1**).

**Figure 3.**
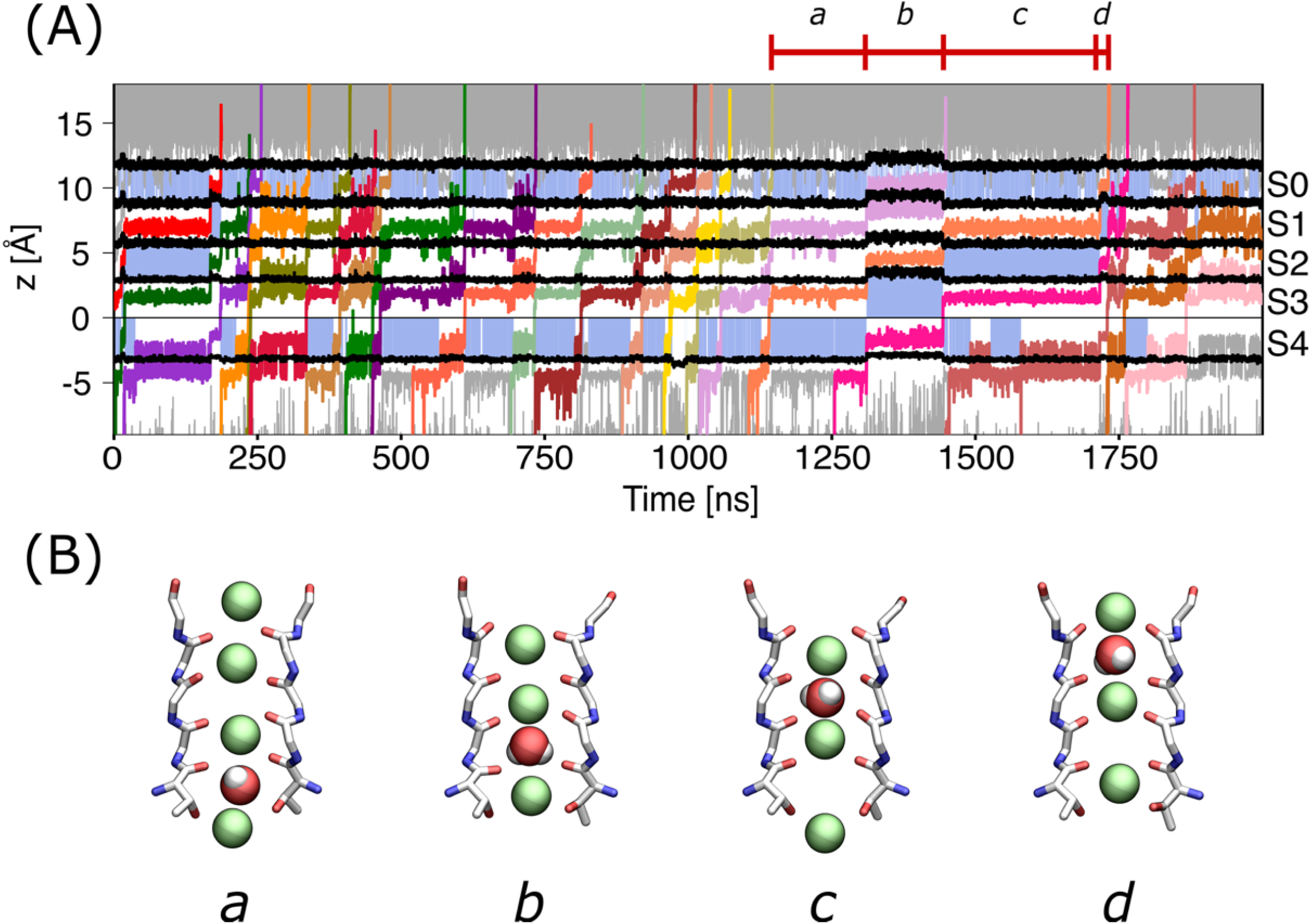
Representative conduction events with hard and soft knock-on mechanism in simulations of MthK with the OPC water model. (A) The axial positions of ions crossing the SF are shown for a representative simulation of MthK with the OPC water model at a membrane potential equal to 200 mV. Different colours are used to represent potassium ions crossing the selectivity filter, while other potassium ions in the system are shown as grey lines. Black lines indicate the boundaries between binding sites S0 to S4, defined by the average positions along the channel axis of the delimiting oxygen atoms. Sky-blue shading denotes frames in which binding sites S0-S4 are occupied by water molecules. (B) Structural representation of the SF during a soft knock-on event. Two opposing subunits of the SF of the MthK channel are shown as sticks, while potassium ions and the water molecule are displayed as van der Waals spheres. States a-d correspond to typical configurations adopted by the SF along a soft knock-on event taken from the timeframes highlighted in panel A.

In order to get a more intuitive and quantitatively accurate description of conduction events, the simulated MD trajectories using the OPC water model were discretized according to the occupancy of the intracellular cavity and SF by ions and water molecules, and then used to estimate MSMs. A schematic representation of the MSM at a membrane potential of 200 mV for MthK and KcsA is shown in **Figure 4**. The size of the nodes is proportional to the probability of the corresponding microscopic states of the MSM, while the colour is based on the presence of water molecules in binding sites S2 and S3, purple indicates the absence of water molecules from both sites, orange indicates the presence of a water molecule in S2, and green indicates the presence of a water molecule in S3. No state was observed with water molecules both in S2 and S3. The width of the edges is proportional to the probability of the corresponding state transition, and the nodes are clustered on the base on these connections, in other words, nodes that are more likely to interconvert are placed closer than those that are not directly connected. These visual representations of the MSMs clearly reveal a clustering of the microscopic states based on the presence of water molecules at the central sites of the SF. In both channels, the water-depleted states at sites S2 and S3 are prevailing, further supporting the dominant role of the hard knock-on mechanism in ion conduction, even when using the OPC water model. Notably, the most populated state differs between the two channels, being wK-KwK in MthK and K-KK-w in KcsA. The latter is consistent with previous studies of ion conduction in KcsA performed with the TIP3P water model.(*16*) More generally, aside from the K-KK-K state, which displays nearly equal populations in both systems, the graphs reveals distinct patterns of ion conduction, extending previous observations of channel-specific ion conduction pathways to channels sharing the same signature sequence in the SF.(*16*) Regarding the soft knock-on-competent states, configurations with a water molecule in S2 are more likely than those with water in S3.

However, the most striking feature emerging from these analyses is the relative paucity of alternating wKwKwK/KwKwKw states predicted by the conventional soft knock-on model mechanism. In particular, in MthK, the KwKwKw state is virtually absent (probability below 10^-3^), while in KcsA, the wKwKwK KwKwKw transitions occur only rarely, suggesting a more complex mechanism of water co-transport than previously proposed.

**Figure 4.**
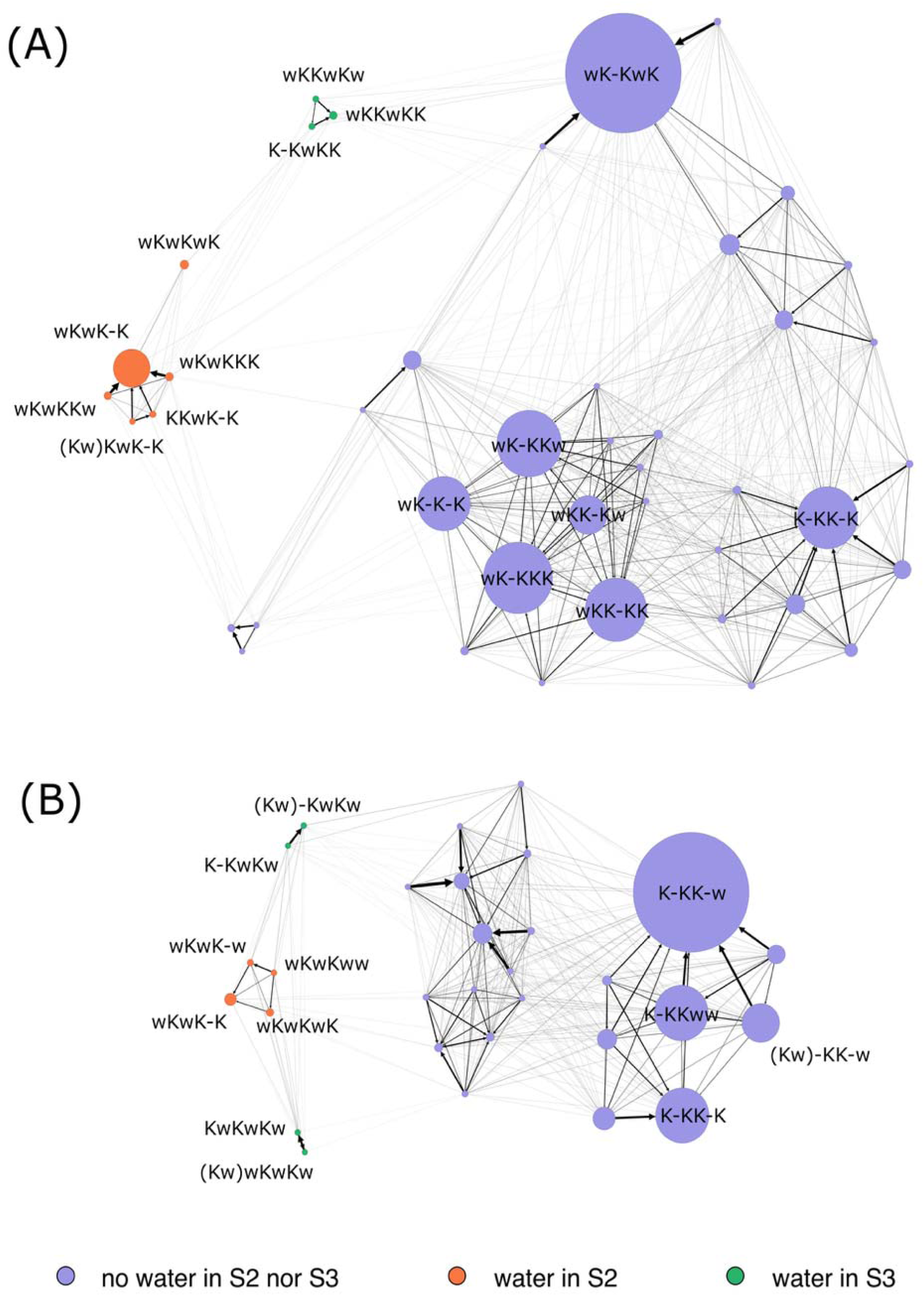
Graph representation of the MSM for MthK (A) and KcsA (B) in simulations with the OPC water model and membrane potential equal to 200 mV. The size of the nodes is proportional to the steady-state probability, and the colour is associated with the presence of water molecules in S2 (orange), S3 (green), or their absence at both sites (purple). Nodes corresponding to states with probabilities below 10^-3^ are not shown. The width of the edges is proportional to the probability of the corresponding transition in the MSM estimated at sampling time of 1 ns. The distinct states are labelled according to the presence of water and potassium in S0, S1, S2, S3, S4, and C, respectively. A dash represents vacant sites. A water/potassium double occupancy occasionally occurring in site S0 is denoted by parentheses. For clarity, only the most relevant states are explicitly labelled

The partitioning between states with and without water molecules in S2 and S3 is confirmed by spectral analysis of the transition matrix of the MSMs. The highest eigenvector of the transition matrix with a module strictly lower than 1 identifies the direction of slowest dynamics.. When the microscopic states of the MSMs are projected along this eigenvector, the resulting two macroscopic states are differentiated by the presence or absence of water molecules in the central sites of the SF for both channels, at all simulated membrane potentials (**Figure 5** and **S1**, **S2**). The most probable macroscopic state is always the one without water molecules in S2/S3, while the minor state consistently shows a significantly higher probability of water being present in these sites. Remarkably, the timescale of the slowest dynamics, as defined by the first eigenvalue less than one, is typically much longer than the next highest timescale (**Figure S3**). This gap between the first and the second timescales indicates that the entrance or exit of water molecules in S2/S3 is the rate limiting step in the simulated systems. In practice, when using the OPC water model, the ion channels switch between two metastable states: one without water molecules in S2/S3, where conduction occurs via the hard knock-on mechanism, and one with water molecules in S2/S3, where water is co-transported with ions, as expected in the soft knock-on mechanism. The separation of the first timescale from the following ones is significantly smaller in the MthK channel at 400 mV compared to other simulated conditions, indicating that in this case, the entrance or exit of water molecules in the SF occurs on a timescale comparable to other events involved in ion conduction.

**Figure 5.**
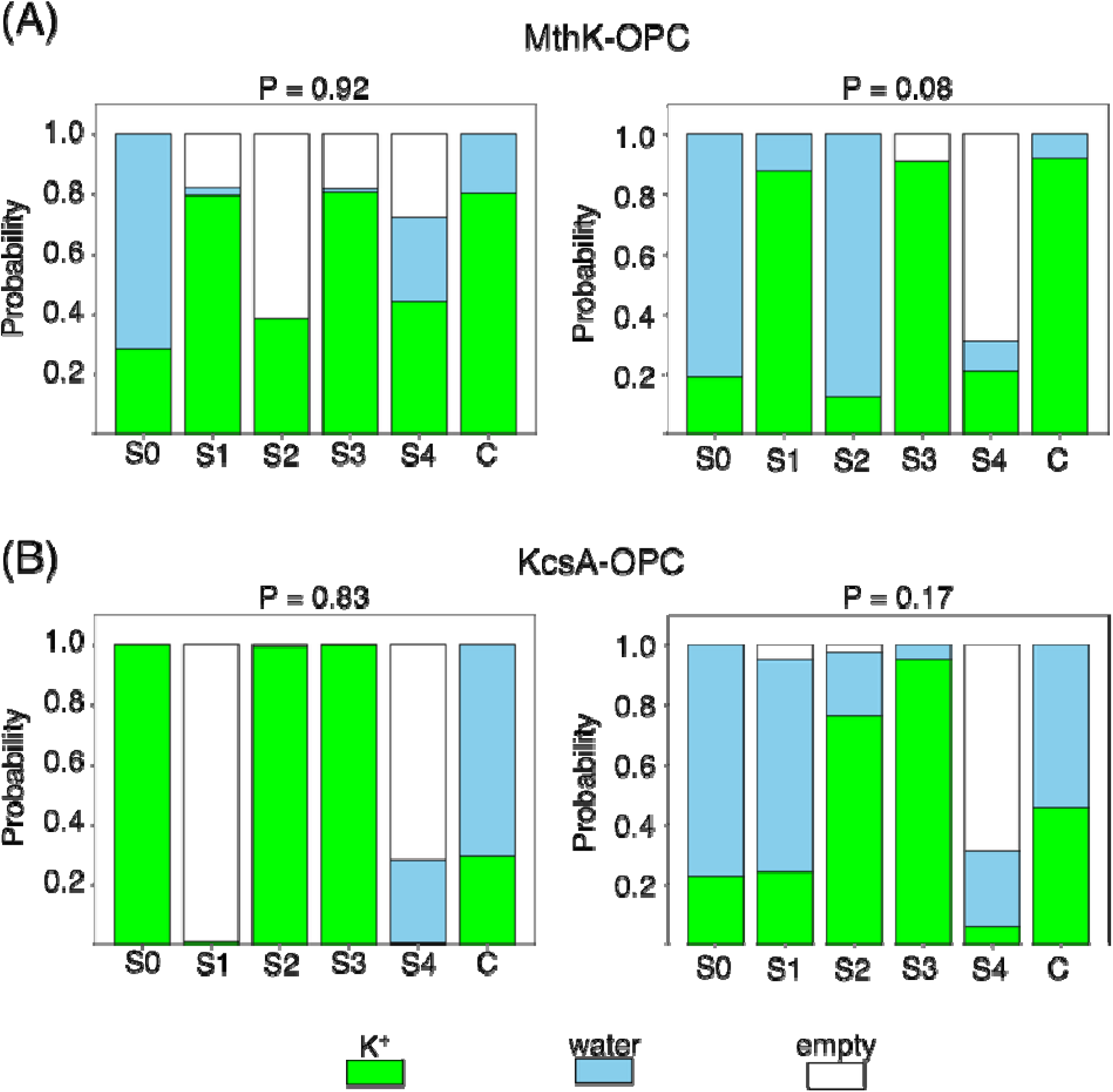
Average state of the SF in a two state models that separates slowest converting microstates in (A) MthK and (B) KcsA. The two macrostates were defined according to the sign of their projection along the eigenvector of the MSM corresponding to the slowest timescale. The average state of the SF was computed as the weighted average of the probabilities of the binding sites being empty, occupied by water, or occupied by potassium ions within two clusters. The cumulative probability of the two clusters is reported at the top. Data refers to the MSM estimated at sampling period of 1 ns for simulations of MthK/KcsA with the OPC water model at a membrane potential of 200 mV.

The MSMs offer the possibility of investigating how the system switches between the two metastable states characterized by the presence or absence of water molecules in S2/S3. The microscopic states that lead to the entrance of water molecules in S2/S3 are characterized by a distribution in which, on average, S2 is empty, S3 is occupied by an ion, S4 by a water molecule, and an ion is present in the intracellular cavity (**Figure 5** and **S4**, **S5**). In practice, the transition from hard to soft knock-on conduction is typically triggered by the formation of a gap in S2, which draws in the ion-water-ion chain inward from the intracellular side of the SF. The opposite event, *i.e.* the switch from soft to hard knock-on conduction, is usually triggered by a state with water in S0, ion in S1, water in S2, and ion in S3, suggesting that exit of the water molecule from S2 toward the extracellular side causes dehydration of the central sites of the SF. This behaviour is consistently observed across both channels and membrane potentials.

**Figure 6.**
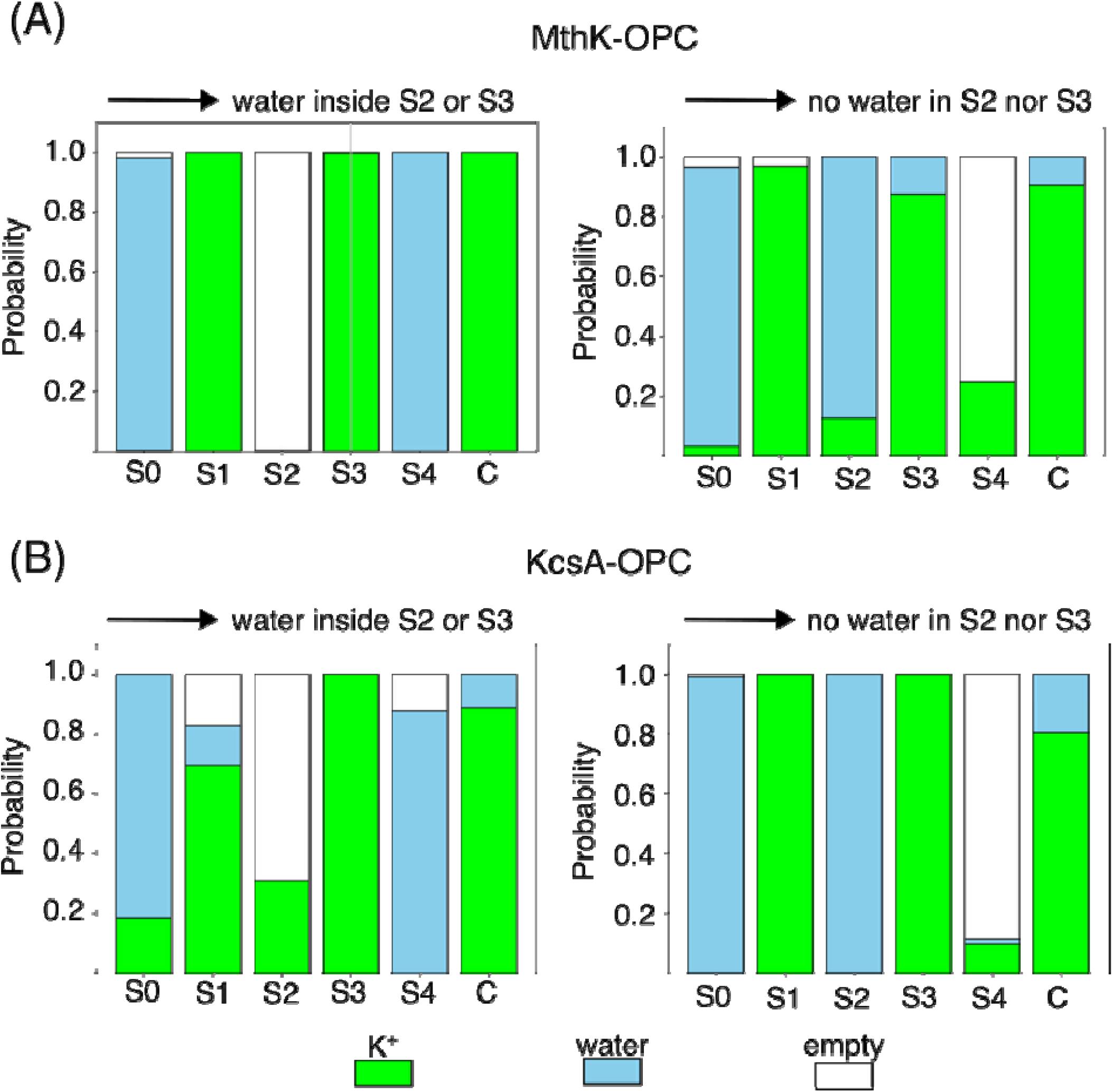
Average state of the SF that precedes the entrance of water molecules in S2 or S3 (left) or the depletion of S2 and S3 from water molecules (right). The average state of the SF that leads to the entrance of water molecules in S2 or S3 was calculated considering all the microscopic states of the MSM with no water in S2 or S3 (*source*) that are connected to any microscopic state with water molecules in S2 or S3 (*sink*). The weighted average probabilities of the binding site being empty, occupied by water, or occupied by potassium ions were then calculated. The same method, switching *source* and *sink*, was used to calculate the average state of the SF that leads to the depletion of water from S2 and S3. Data refer to simulations of the MthK/KcsA channels with the OPC water model at a membrane potential of 200 mV.

## Discussion

The analyses of the MD trajectories we presented here prove that, when simulations of potassium channels adopt the OPC water model, in combination with the protein force-field AMBER19SB and the 12-6-4 Sengupta et al. model of ions, both the hard knock-on and the soft knock-on mechanisms of conduction are possible. In the long-standing argument between these two conduction mechanisms, to the best of our knowledge, this is the first time that soft knock-on is observed with significant probability in MD simulations, and that reversible transitions between hard and soft knock-on are reported. The hard knock-on mechanism was initially presented as a possible alternative to what was at the time the accepted mechanism of conduction that involved water-ion co-permeation, and that was simply known as knock-on conduction.(*30*) The prevalence of hard knock-on over soft knock-on later emerged in numerous reports of MD simulations where drifting potentials were used to force ion conduction, and where hard knock-on was largely the most common, if not the unique, mechanism of conduction.(*14*) Subsequently, X-ray crystallography in the presence of electric field stimulation provided electron densities of potassium channels that are reminiscent of the steps observed in MD simulations of conduction events with the hard knock-on mechanism.(*26*) Despite these data, the debate about the atomic details of conduction in potassium channels, and primarily the possible involvement of water molecules, is still not settled. Most importantly, streaming potential measurements demonstrate the co-permeation of water with ions, which is clearly not compatible with hard knock-on conduction.(*66–68*) An immediate possibility to reconcile streaming potential experiments with the literature supporting hard knock-on conduction is that both conduction mechanisms coexist. This hypothesis of multiple mechanisms of conduction being possible in potassium channels resonates with our previous observations of differences in conduction strategies among potassium channels, and experimental conditions.(*16*) Under this perspective, it is interesting to note the coexistence of hard and soft knock-on observed here in MD simulations with the OPC water model.

The transition between hard and soft knock-on conduction was reversible in both channels, MthK and KcsA, and at all simulated membrane potentials. These reversible transitions are in stark contrast to previous observations in molecular dynamics simulations using the TIP3P water model. When TIP3P is combined with the CHARMM36m force-field, the entrance of water molecules in the central binding sites of the SF triggers a series of structural changes that ultimately leads to a non-conductive state of the channel. Instead, when TIP3P is combined with the AMBER14SB force-field, water molecules were never observed to reach sites S2/S3 in microsecond trajectories.(*30*) In cases were the systems are prepared with water molecules in S2 or S3, the water is eventually expelled, and afterwards pure hard knock-on conduction is established. The reversible transitions observed here in simulations with OPC are necessary for the coexistence of hard and soft knock-on.

The two simulation setups considered in this work differ in several important aspects beyond the water model, including the protein force-field and the ion parameters. Identifying which specific factor is responsible for the differences that we observed remains an interesting question, which might provide important clues for further force-field development. The aim of our analyses was to evaluate whether simulations of potassium channels with OPC water models yield results significantly different than previous reports in the literature employing other water models, mainly TIP3P. The combination of OPC with a force-field, such as AMBER14SB, developed for TIP3P, would not answer our question. Instead, it was natural to combine the OPC water model, with a new generation protein force-field which proved better performances with such water model, and specifically parametrized ion parameters, which is also how simulations with OPC are usually performed. The results clearly show that this combination of parameters produces conduction mechanisms that differ markedly from the ones observed with other parameter sets, including amber- and charmm-based force-fields. While previous MD simulations, mainly using the TIP3P water model, practically ruled out soft knock-on as a possible mechanism of conduction, our findings bring it back into the discussion.

## Author Contributions

The manuscript was written through contributions of all authors. All authors have given approval to the final version of the manuscript.

## Acknowledgments

We acknowledge CINECA for awarding access to computational resources through the ISCRA Initiative (grant numbers HP10B597KB and HP10B5IPGG). Riccardo Ocello is gratefully acknowledged for useful discussion. We acknowledge EuroHPC for awarding access to computational resources in several of the European platforms including LUMI (CSC, Finland), Marenostrum (BSC, Spain) and Leonardo (CINECA, Italy).

## Notes

Configuration files, atomic models, discretized MD trajectories, and the python code used for the analyses of the MSM are available at the github repository: https://github.com/sfurini/kchannels_water_models

## Abbreviations

MD: Molecular Dynamics
SF: Selectivity Filter

