## Supplementary Material for "Role of Water Models in Simulations of Ion Conduction in Potassium Channels"

*** Corresponding authors**

l

**Table S1. Regions used to discretize the MD trajectories of KcsA.** Ions and water molecules were considered inside S0-S4, and C, when within these boundaries. Extracellular and Intracellular boundaries were computed as centre of mass of the corresponding group of atoms.

|  | **Extracellular Boundary** | **Intracellular Boundary** | **Maximum distance from the channel axis [Å]** |
| --- | --- | --- | --- |
| S0 | backbone oxygens of Gly79 | backbone oxygens of Tyr78 | 4 |
| S1 | backbone oxygens of Tyr78 | backbone oxygens of Gly77 | 4 |
| S2 | backbone oxygens of Gly77 | backbone oxygens of Val76 | 4 |
| S3 | backbone oxygens of Val76 | backbone oxygens of Thr75 | 4 |
| S4 | backbone oxygens of Thr75 | hydroxyl oxygens of Thr75 | 4 |
| C | hydroxyl oxygens of Thr75 | residues Thr107 | 8 |

**Table S2. Regions used to discretize the MD trajectories of MthK.** Ions and water molecules were considered inside S0-S4, and C, when within these boundaries. Extracellular and Intracellular boundaries were computed as centre of mass of the corresponding group of atoms.

|  | **Extracellular Boundary** | **Intracellular Boundary** | **Maximum distance from the channel axis [Å]** |
| --- | --- | --- | --- |
| S0 | backbone oxygens of Gly63 | backbone oxygens of Tyr62 | 4 |
| S1 | backbone oxygens of Tyr62 | backbone oxygens of Gly61 | 4 |
| S2 | backbone oxygens of Gly61 | backbone oxygens of Val60 | 4 |
| S3 | backbone oxygens of Val60 | backbone oxygens of Thr59 | 4 |
| S4 | backbone oxygens of Thr59 | hydroxyl oxygens of Thr59 | 4 |
| C | hydroxyl oxygens of Thr59 | residues Glu92 | 8 |

**Table S3. List of MD simulations and corresponding conduction events.** Conduction events were considered as soft knock-on when a water molecules moves across the SF.

| **Channel** | **Water model** | **Membrane Potential [mV]** | **MD simulations** | **Hard knock-on conduction events** | **Soft knock-on conduction events** |
| --- | --- | --- | --- | --- | --- |
| KcsA | TIP3P | 100 | 7 x 4.0 µs | 60 | 0 |
|  |  | 200 | 8 x 4.0 µs | 346 | 0 |
|  |  | 400 | 8 x 1.0 µs | 436 | 0 |
| KcsA | OPC | 100 | 4 x 3.0 µs and 4 x 2.0 µs | 16 | 11 |
|  |  | 200 | 4 x 3.0 µs and 4 x 2.0 µs | 52 | 10 |
|  |  | 400 | 8 x 2 µs | 234 | 42 |
| MthK | TIP3P | 100 | 8 x 2 µs | 27 | 0 |
|  |  | 200 | 4 x 2.0 µs and 4 x 1.0 µs | 150 | 0 |
|  |  | 400 | 4 x 1.5 µs and 4 x 1.0 µs | 474 | 0 |
| MthK | OPC | 100 | 8 x 2 µs | 34 | 4 |
|  |  | 200 | 4 x 3.0 µs and 4 x 2.0 µs | 113 | 6 |
|  |  | 400 | 8 x 2 µs | 173 | 37 |

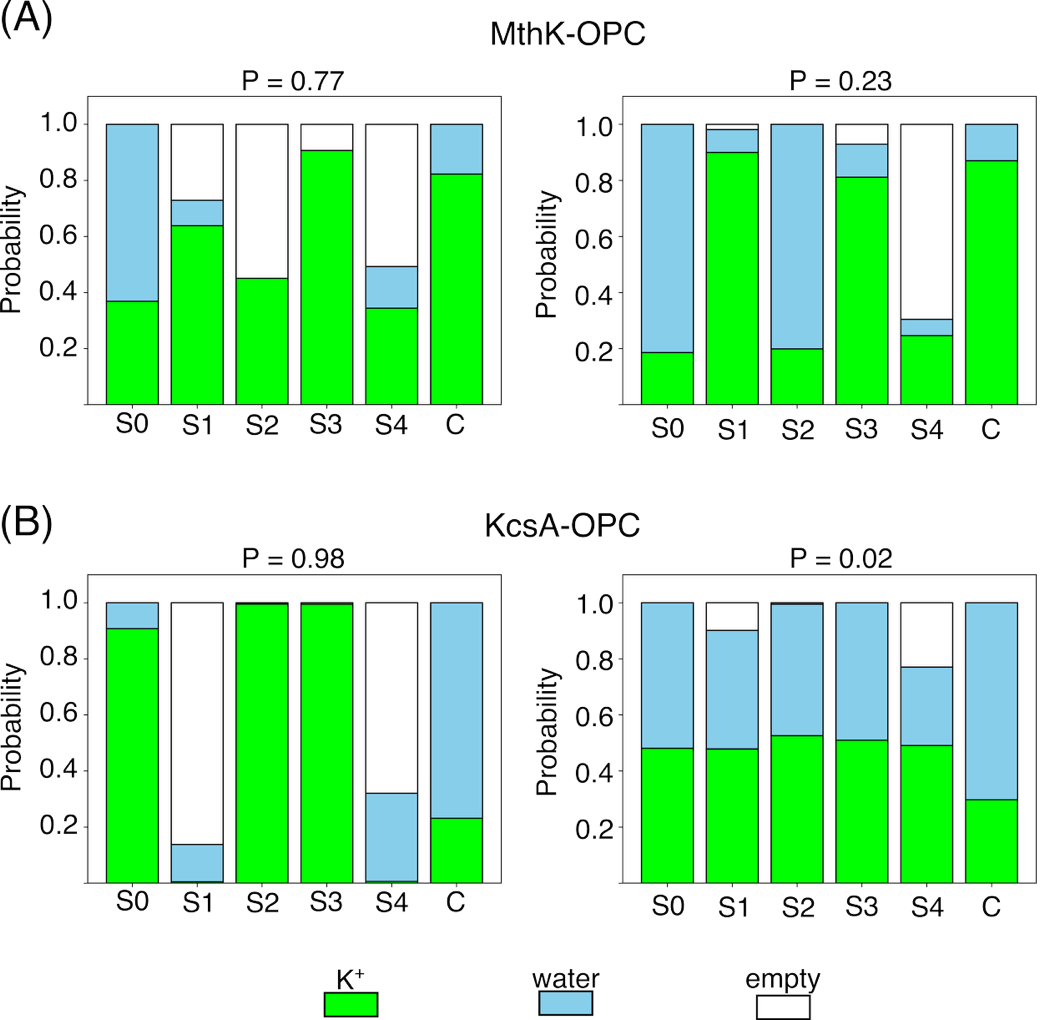

**Figure S1. Average state of the SF in a two state models that separates slowest converting microstates in MthK (A) and KcsA (B) at 100 mV.** The two macrostates were defined according to the sign of their projection along the eigenvector of the MSM corresponding to the slowest timescale. The average state of the SF was computed as the weighted average of the probability of the binding sites being empty, occupied by water, or occupied by potassium ions, respectively in the two clusters. The cumulative probability of the two clusters is reported at the top. Data refers to the MSM estimated at sampling period of 1 ns for simulations of MthK/KcsA with the OPC water model at membrane potential equal to 100 mV.

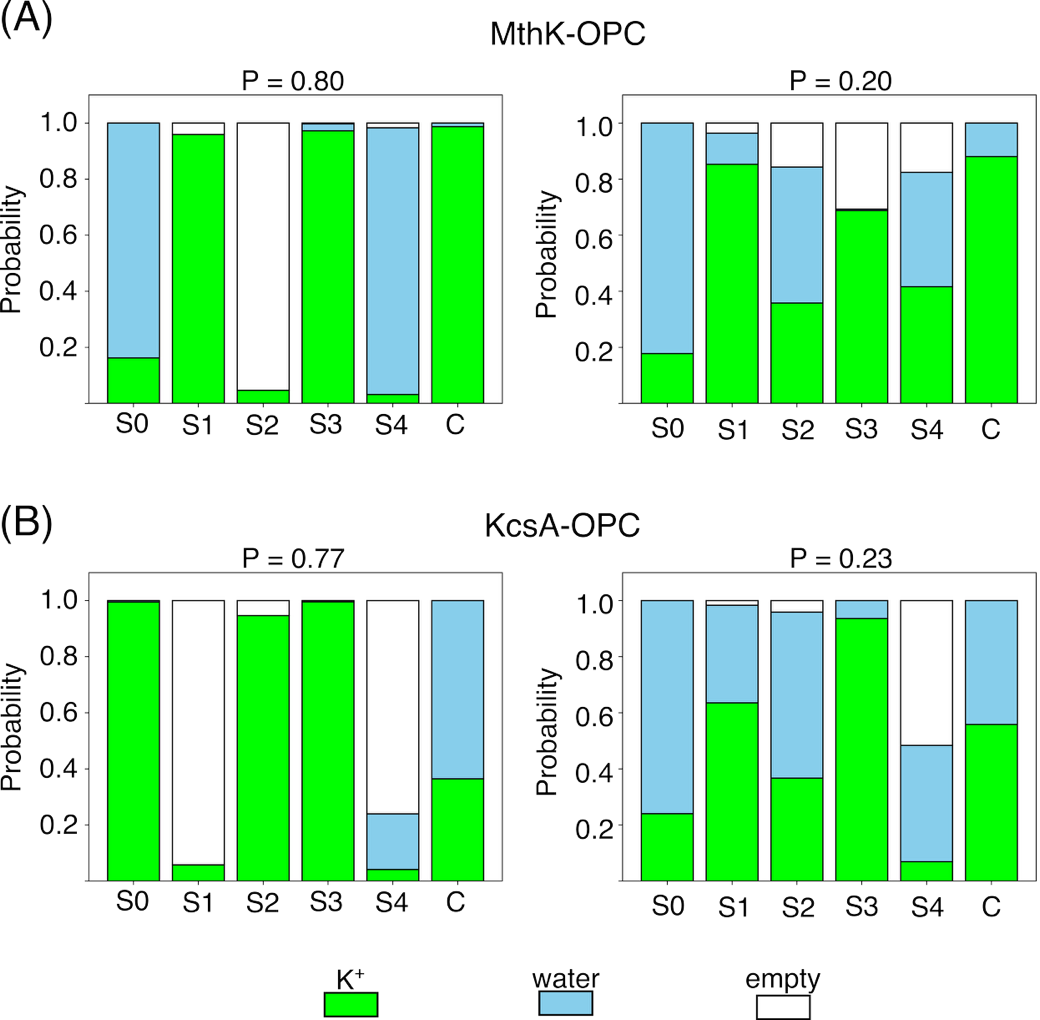

**Figure S2. Average state of the SF in a two state models that separates slowest converting microstates in MthK (A) and KcsA (B) at 100 mV.** The two macrostates were defined according to the sign of their projection along the eigenvector of the MSM corresponding to the slowest timescale. The average state of the SF was computed as the weighted average of the probability of the binding sites being empty, occupied by water, or occupied by potassium ions, respectively in the two clusters. The cumulative probability of the two clusters is reported at the top. Data refers to the MSM estimated at sampling period of 1 ns for simulations of MthK/KcsA with the OPC water model at membrane potential equal to 100 mV.

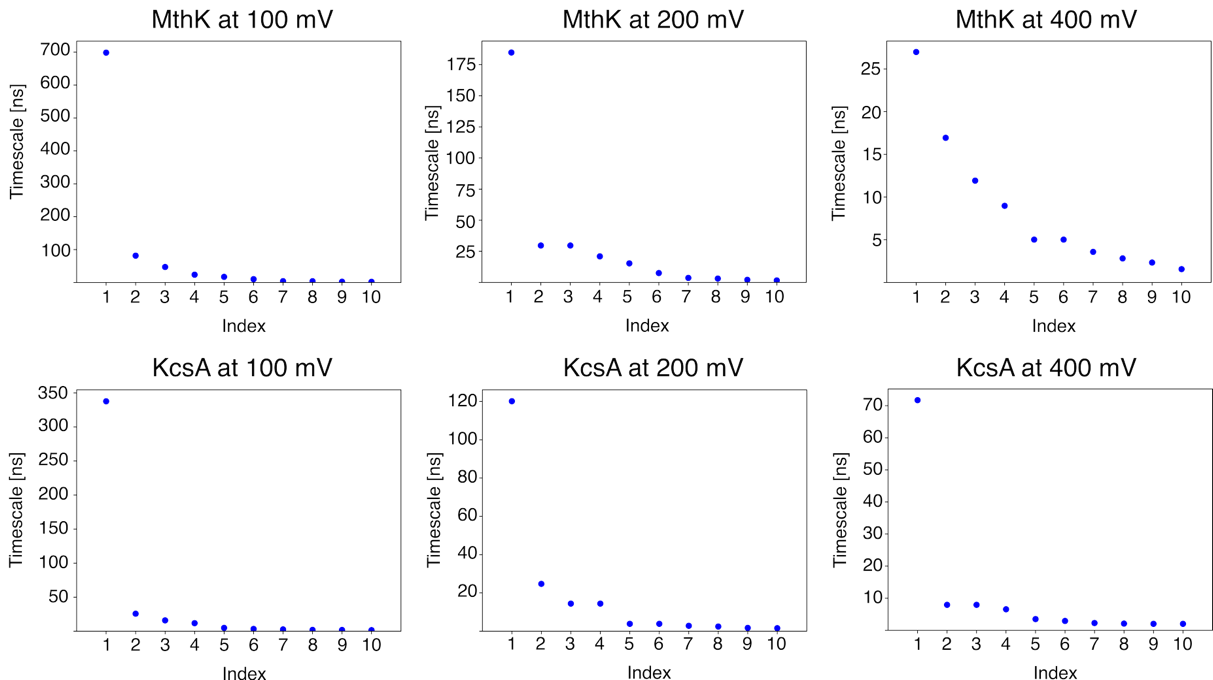

**Figure S3.** Relaxation times estimated from the transition matrix of the MSMs.

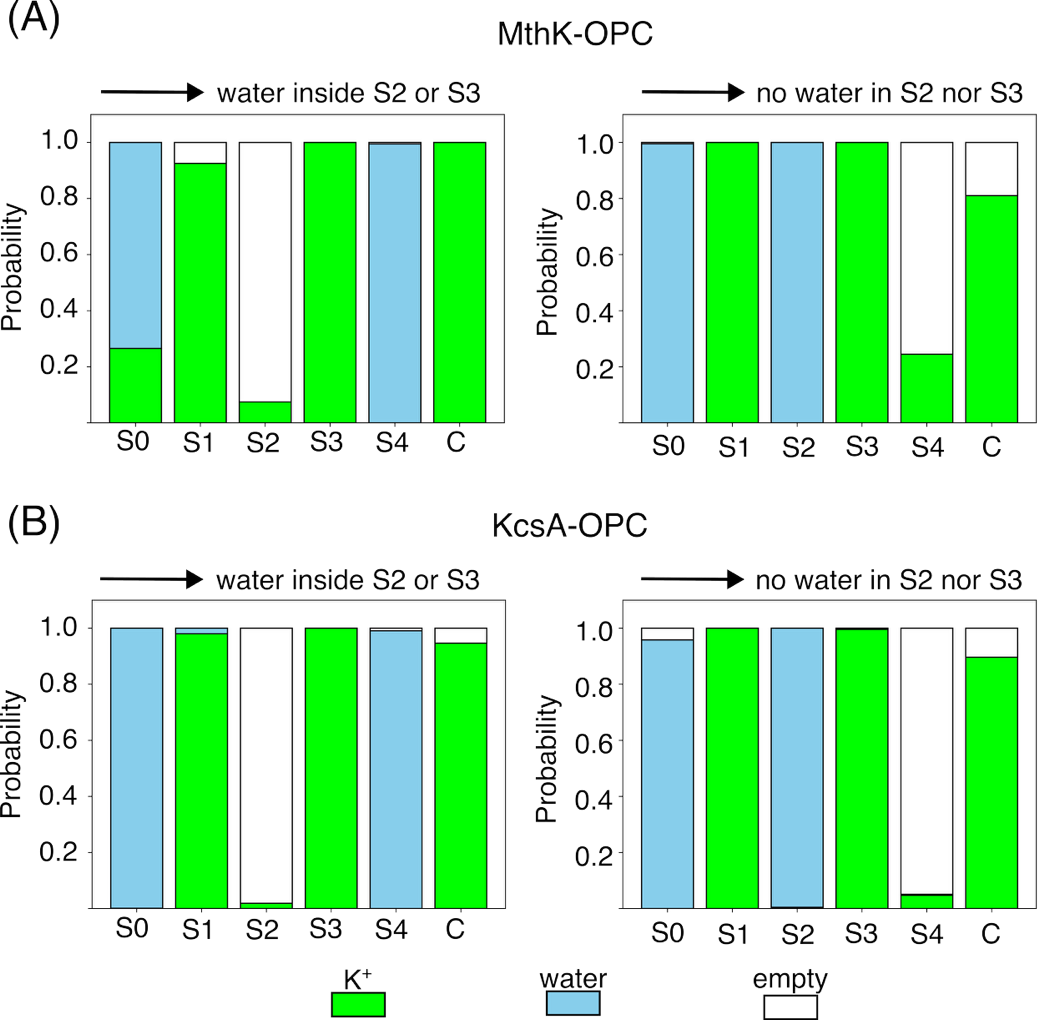

**Figure S4. Average state of the SF that precedes the entrance of water molecules in S2 or S3 (left) or the depletion of S2 and S3 from water molecules (right).** The average state of the SF that leads to the entrance of water molecules in S2 or S3 was calculated considering all the microscopic states of the MSM with no water in S2 nor S3 (*source*) that are connected to any microscopic state with water molecules in S2 or S2 (*sink*). Then, the weighted average of the probability of the binding site being empty, occupied by water, or occupied by potassium ions were calculated. The same method, switching *source* with *sink*, was used to calculate the average state of the SF that leads to the depletion of S2 and S3 from water molecules. Data refers to simulations of the MthK/KcsA channel with the OPC water model at a membrane potential of 100 mV.

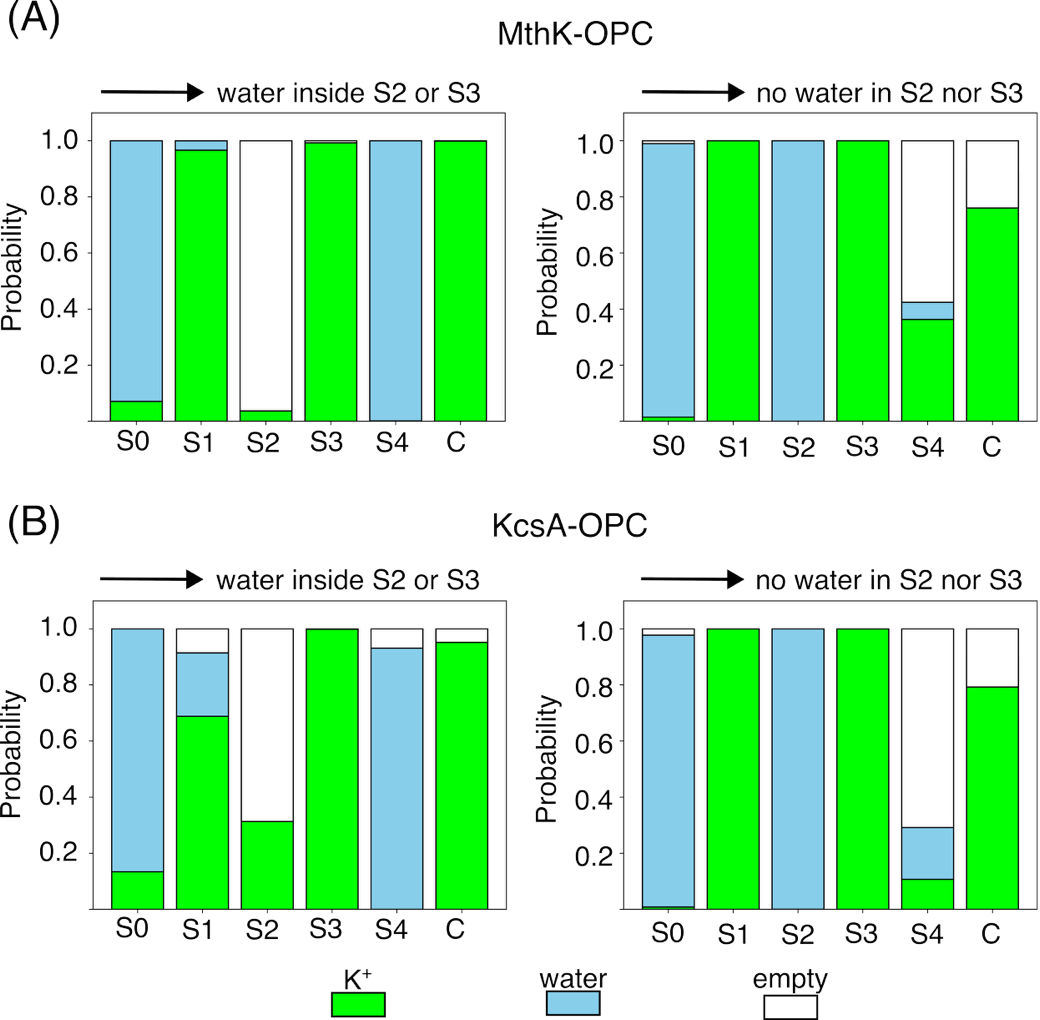

**Figure S5. Average state of the SF that precedes the entrance of water molecules in S2 or S3 (left) or the depletion of S2 and S3 from water molecules (right).** The average state of the SF that leads to the entrance of water molecules in S2 or S3 was calculated considering all the microscopic states of the MSM with no water in S2 nor S3 (*source*) that are connected to any microscopic state with water molecules in S2 or S2 (*sink*). Then, the weighted average of the probability of the binding site being empty, occupied by water, or occupied by potassium ions were calculated. The same method, switching *source* with *sink*, was used to calculate the average state of the SF that leads to the depletion of S2 and S3 from water molecules. Data refers to simulations of the MthK/KcsA channel with the OPC water model at a membrane potential of 400 mV.
